# Aquifer microbial communities are highly variable in response to thermal arsenic mobilization

**DOI:** 10.1101/2024.10.03.616553

**Authors:** Molly Chen, Daniel S. Grégoire, Pascale St-Germain, Carolina Berdugo-Clavijo, Laura A. Hug

## Abstract

Thermal recovery technologies for *in-situ* bitumen extraction can result in the heating of surrounding aquifers, potentially mobilizing arsenic naturally present in the sediments to the groundwater. The relative toxicity of dissolved arsenic is related to its speciation, with As(V) being less toxic than As(III). Microorganisms have various mechanisms of arsenic resistance, including efflux and methylation. Microorganisms may also perform reduction/oxidation of As(V)/As(III) as part of their detoxification and/or metabolic pathways. We characterized the microbial communities along two aquifer transects associated with thermally mobilized arsenic near Northeastern Alberta oil sands deposits. 16S rRNA amplicons and metagenomic sequencing data of biomass from filtered groundwater indicated major changes in the dominant taxa between wells, especially those currently experiencing elevated arsenic concentrations. Annotation of arsenic-related genes indicated that efflux pumps (*arsB, acr3*), intracellular reduction (*arsC*) and methylation (*arsM*) genes were widespread amongst community members but comparatively few organisms encoded genes for arsenic respiratory reductases (*arrA*) and oxidases (*arxA, aioA*). While this indicates that microbes have the capacity to exacerbate arsenic toxicity by increasing the relative concentration of As(III), some populations of iron oxidizing and sulfate reducing bacteria (including novel *Gallionella* and Thermodesulfovibrionia populations) show potential for indirect bioremediation through formation of insoluble iron/sulfide minerals which adsorb or coprecipitate arsenic. An unusually high proportional abundance of a single Paceibacteria population that lacked arsenic resistance genes was identified in one high-arsenic well, and we discuss hypotheses for its ability to persist. Overall, this study describes how aquifer microbial communities respond to thermal and arsenic plumes, and predicts potential contributions of microbes to arsenic biogeochemical cycling under this disturbance.

## 1. INTRODUCTION

Global health concerns regarding arsenic exposure are mainly tied to the contamination of groundwater used for drinking and agriculture (Mukherjee et al., 2006). Arsenic can be mobilized to groundwaters through natural sources, including hydrothermal activity and mineral weathering processes (reviewed in Masuda, 2018) and anthropogenic activities, including natural resource extraction (e.g., mining) and rice agriculture. As a result, high concentrations of arsenic (above the WHO guideline of 0.01 mg/L for safe drinking water) are present in aquifers in many countries (World Health Organization, 2022). Chronic arsenic exposure is linked to cancer and skin lesions, among other adverse effects (World Health Organization, 2022). It is estimated that 94-220 million individuals are at risk of drinking arsenic-contaminated water (Podgorski & Berg, 2020).

The bioavailability and toxicity of inorganic arsenic is controlled by abiotic and biotic pathways that affect arsenic’s redox state. Arsenic forms oxyanions in solution, as either the trivalent (As^(III)^(OH)_3_) or pentavalent (H_2_As^(V)^O_4_^-^ or HAs^(V)^O_4_^2-^) species. As(III) is considered more toxic than As(V) (Akter et al., 2005), and is generally more mobile in natural environments due to its lower adsorption potential to minerals and sediments (Smedley & Kinniburgh, 2002). Arsenic speciation in groundwater is primarily determined by pH and redox potential: Under acidic/circumneutral pH and reducing conditions, As(III) is the major dissolved species, while As(V) is predominant in high pH (>8.5) and oxidizing environments (Smedley & Kinniburgh, 2002). These variables also control whether arsenic is mobilized through reductive dissolution pathways or immobilized through precipitation and adsorption onto nearby sediments.

Microorganisms play a major role in arsenic biogeochemical cycling through direct and indirect mechanisms. Direct mechanisms involve the expression of genes that specifically interact with arsenic for detoxification or contributing to cellular metabolism (reviewed in Crognale et al., 2017; Zhu et al., 2017). The most common intracellular detoxification pathway involves the *ars* and/or *acr* operons, which are widely distributed in bacteria and archaea (Fu et al., 2009; Silver & Phung, 2005). The ArsB/Acr3 efflux pump expels intracellular As(III) and is coupled to the activity of the intracellular reductase, ArsC, which reduces As(V) to As(III) prior to efflux (Mukhopadhyay & Rosen, 2002). Methylation of As(III) (by ArsM) to methylarsenite/methylarsenic acid (MAs(III)) is a microbial method of producing antibiotics to inhibit competitors, as MAs(III) is more toxic than the inorganic form (Chen & Rosen, 2020). However, As(III) methylation can also confer arsenic resistance if MAs(III) is oxidized to MAs(V), which is less toxic than either inorganic form of arsenic and favoured in the presence of oxygen (Chen & Rosen, 2020).

Many bacteria and archaea are also able to obtain energy by using As(III) as an electron donor or As(V) as a terminal electron acceptor. As(III) oxidases include AioAB (present in various heterotrophs and chemolithoautotrophs) and ArxA (primarily in Gammaproteobacteria) (Shi et al., 2020). The dissimilatory (or respiratory) As(V) reductase ArrAB differs from ArsC, in that it is membrane-bound and reduces extracellular As(V) to As(III) as part of an electron transport chain (Basu et al., 2010; Malasarn et al., 2004), rather than for detoxification. Dissimilatory As(V) reduction is common in many microorganisms found in groundwater ecosystems (Crognale et al., 2017; Escudero et al., 2013).

Indirect mechanisms of arsenic biogeochemical cycling are connected to the iron and sulfur cycles, among others (reviewed in Zhu et al., 2017). The distribution and speciation of iron and arsenic in sediments are highly correlated, as arsenic is strongly sorbed to Fe(III) (oxy)hydroxides and a constituent of many Fe-bearing minerals, such as arsenopyrite (Fe(S,As)_2_) (Smedley & Kinniburgh, 2002). Fe(III) reduction and Fe(II) oxidation by microbes can lead to the dissolution or precipitation of these minerals, depending on pH and redox conditions, which controls the mobilization of arsenic (Zhu et al., 2017). Another important mechanism for attenuation of arsenic is dependent on sulfur cycling. Sulfate-reducing bacteria (SRBs) produce biogenic sulfide, leading to the subsequent formation of arsenopyrite and other arsenic-bearing sulfide minerals (*e.g.,* realgar, AsS; orpiment, As_2_S_3_) (Newman et al., 1997; Rittle et al., 1995). Although biological arsenic cycling has been well-studied in the context of natural resource extraction in mining and mine tailings (Akter et al., 2005; Pakostova et al., 2023; Zhuang et al., 2023), how microbial activities impact the fate of arsenic in petroleum extraction operations remains understudied.

We are addressing this gap by examining the biogeochemical cycling of arsenic at a model petroleum extraction site in Northern Alberta. This site uses steam to extract the bitumen from heavy oil deposits. During this process, high temperature steam (∼300 °C) is injected to the reservoir to liquify the bitumen and subsequently pump to surface facilities. This results in conductive heating of the surrounding sediments and subsequent mobilization of arsenic naturally present in sediments into groundwater (Moncur, 2018). Desorption from metal oxides is the dominant mechanism of thermal arsenic mobilization (Craig et al., 2021), but reductive dissolution of arsenic-bearing minerals also accelerates under higher temperatures (Johnston et al., 2020; Weber et al., 2010). Experiments on sediment cores in close proximity to steam injection wells showed that heating of sediments greatly increased arsenic concentrations in porewater, from a baseline of <0.025 mg/L up to a maximum of 5.200 mg/L (Javed & Siddique, 2016; Moncur et al., 2015). Although these studies highlight how abiotic processes associated with the steam injection process impact arsenic concentrations within the local aquifer system (Javed et al., 2014), how this mobilized arsenic affects microbial community dynamics and microbial arsenic cycling pathways remains unexplored.

In this study, we investigate microbial diversity and activities in groundwater aquifers in response to thermally mobilized arsenic from an *in-situ* bitumen extraction operation. A combined 16S rRNA amplicon and metagenomic sequencing approach was used to profile the taxonomic composition of communities and predict potential interactions between individual organisms and arsenic. We describe the contributions of microbes to direct arsenic cycling within the system (via predicted resistance mechanisms and metabolic redox transformations) as well as the biogeochemical cycling of other elements that may influence arsenic mobility (*e.g.*, Fe, S). These findings are discussed in the context of designing effective bioremediation strategies for arsenic-contaminated aquifers.

## 2. MATERIALS AND METHODS

### 2.1 Site properties and sampling

The study sites are located within Northern Alberta, Canada, in an area containing a series of non-saline aquifers in the Quaternary deposits. *In-situ* extraction of bitumen via cyclic steam stimulation (CSS) is ongoing in the bitumen reservoir, located about 350 m below the aquifers (Andriashek, 2003).

Sampling of groundwater took place in August 2021 across two transects associated with steam injection pads (CSS well pads): Site A and Site B. The Site A aquifer is located approximately 80 m below ground and has a water flow rate of ∼100 m/year. The aquifer has experienced historic arsenic plumes from past steam cycles (1990-2010), but these plumes have mostly dispersed. In contrast, Site B’s aquifer is located at a depth of ∼160 m and has a much slower flow rate (< 5 m/year). Site B is currently experiencing slow-moving plumes of arsenic from steaming during 2003-2013.

Water (100 L) was pumped from a series of wells along each transect (five wells along Site A, six wells along Site B), and immediately filtered through a Waterra 0.1-micron filter (CAP600X-0.1). The filters were purged with air and 50 mL RNAlater (Invitrogen AM7021) was added to preserve biomass prior to filters being transported on ice to the University of Waterloo where they were stored at -80 °C.

Geochemical assessments of temperature, pH, Eh, and EC were measured on-site during groundwater filtering. Additional groundwater was analyzed by Bureau Veritas (see Data File S1 for details on analyses) for concentrations of major cations, anions, alkalinity, and dissolved carbon. Speciation data for dissolved iron (Fe^2+^/Fe^3+^) and arsenic (As^3+^/As^5+^) were measured by Brooks Applied Labs (see Data File S1 for details on analyses).

All wells were sampled again in August 2022 following the same protocol, with the exception of the volume of water filtered (increased to a target of 150 L, with 100-160 L filtered depending on whether the filter clogged).

### 2.2 DNA extraction

Waterra filters were thawed at room temperature for 1-2 hours. After purging the RNAlater through the outlet, the filter membrane was rinsed with 50 mL of Longmire (minus SDS) buffer (0.1M Tris-HCl, 0.1M EDTA, 0.01M NaCl) three times, vortexed for 30 s, and the buffer was collected out of the inlet. The combined 150 mL of buffer with resuspended biomass was filtered onto a Stericup vacuum filter (Millipore S2VPU02RE, 0.1 µm pore size), and the filter membrane was cut into several pieces using a sterile scalpel. Approximately one-third of each filter was directly transferred to the QIAGEN DNeasy® PowerSoil® Pro Kit for DNA extraction; the remainder was stored in 5 mL centrifuge tubes at -80 °C.

DNA was eluted in 60 µL of 10mM Tris and stored at -20 °C. 20 µL was used for 16S rRNA amplicon sequencing. Due to low concentrations (< 3 ng/µL) in 6 out of 11 samples, another third of the filters were used for DNA extraction and combined with the remaining DNA from the first extraction for metagenomic sequencing. Final DNA quantities sent for metagenomic sequencing ranged from 35.2 ng – 187.2 ng.

### 2.3 16S rRNA amplicon sequencing & analysis

Samples were PCR-amplified and sequenced by MetagenomBio Life Science Inc. using the Illumina MiSeq machine (v2 reagents). Modified universal primers targeting the V4 region were used for amplification (forward primer 515FB: 5’ GTGYCAGCMGCCGCGGTAA; reverse primer 806RB: 5’ GGACTACNVGGGTWTCTAAT) (Walters et al. 2015).

Processing and analysis of sequencing data was performed within the QIIME2 platform (v.2022.2) (Bolyen et al., 2019). Paired-end reads were filtered, trimmed, merged, and grouped into amplicon sequence variants (ASVs) with DADA2 (Callahan et al., 2016). Taxonomy was assigned based on the SILVA 138 SSU database (Yilmaz et al., 2014). Alpha and beta diversity were computed using the QIIME2 core phylogenetic diversity metrics tool.

Data (including ASV abundance, taxonomy, diversity tables/matrices, and metadata) were exported for further processing in Python v3.8.8 using the pandas v1.4.4 library (McKinney, 2010). Ordinations and statistical analyses were performed using the scikit-bio v0.5.8 package (Bradbury et al., 2022) in Python, except for environmental fitting, which was done using the vegan v2.6-4 library (Oksanen, 2010) in R v4.2 (R Core Team, 2021). Figures were generated using the seaborn (Waskom, 2021) and matplotlib v3.6.0 (Hunter, 2007) libraries.

### 2.4 Metagenomic sequencing, assembly, and binning

Sequencing was performed at The Center for Applied Genomics (TCAG) (Toronto, Ontario), using PCR-amplified library preparation and 2 lanes of an Illumina Novaseq SP flowcell (PE2x150bp). Raw reads were decontaminated using BBduk (Bushnell, n.d.), and sickle v1.33 (Joshi & Fass, 2011) was used to trim/filter low-quality reads.

Reads were assembled using SPAdes v3.15.5 (Bankevich et al., 2012) with the --meta flag for metagenomes. Assembled scaffolds were filtered to a minimum length of 2500 bp using pullseq v1.0.2 (Thomas, 2015).

Read mapping was done with Bowtie2 v2.3.4.1 (Langmead & Salzberg, 2012) to generate coverage information for binning. Three binning algorithms were used: CONCOCT (Alneberg et al., 2014), MetaBAT 2 (Kang et al., 2019), and MaxBin 2.0 (Wu et al., 2016). The dereplication, aggregation, and scoring tool (DAS Tool) (Sieber et al., 2018) was used to select the best set of high-quality non-redundant bins from the candidate bins. CheckM (Parks et al., 2015) was used to filter bins for >70% completeness and <10% contamination as the final set of high-quality bins.

### 2.5 Taxonomic placement and gene annotation of MAGs

Taxonomy of MAGs was assigned using the Genome Taxonomy Database Toolkit (GTDB-Tk v2.1.0) (Chaumeil et al., 2020). A phylogenetic tree was generated using GToTree v1.6 (M. D. Lee, 2019), and visualized using iToL (Letunic & Bork, 2021).

Annotation of arsenic-related genes was performed with the meta-arsenic toolkit (Dunivin et al., 2019), which incorporates hidden Markov models (HMMs) and BLAST database information to identify the following arsenic resistance and respiratory genes: *acr3*, *aioA*, *arsB*, *arsC* (grx), *arsC* (trx), *arsD*, *arsM*, *arrA*, and *arxA*. FeGenie (Garber et al., 2020) was used to annotate iron-related genes, also using HMM databases. Distilled and Refined Annotation of Metabolism (DRAM v1.5.0) (Shaffer et al., 2020) was used to profile the overall metabolism of MAGs, with annotations labeled by Pfam (El-Gebali et al., 2019) and KEGG (Kanehisa et al., 2017) identifiers. Sulfur-related genes were filtered from the resulting output for prediction of MAGs with sulfur-cycling capacities.

All data from meta-arsenic, FeGenie, and DRAM were exported to Python (pandas library). Arsenic-, iron-, and sulfur-related gene frequencies were aggregated into pathways (*e.g.*, oxidation, reduction), expressed as a binary (indicating presence/absence), and then normalized to MAG % relative abundance as calculated using average scaffold coverage data as a proxy for the prevalence of pathways in each sample. Figures were generated using matplotlib.

### 2.6 Prediction of host-symbiont interactions

To identify potential hosts for the abundant Paceibacteria MAG CLB2-064 population within well B-02, we performed a co-occurrence analysis using the R package propr v2.1.2 (Quinn et al., 2017), which calculates proportionality of compositional data (such as MAG abundances). Bowtie2 coverage information, as a proxy for abundance, for all scaffolds >10 kb from the well B-02 metagenome was compared to the average coverage of CLB2-064 scaffolds. A cutoff of ρ = 0.9 was used to determine significant proportionality between scaffolds, which corresponded to a false discovery rate (FDR) of 0.047. An additional analysis was performed between Paceibacteria CLB2-064 abundance and the abundance of all high-quality bins within B-02. In this case, proportionality was assessed between average scaffold coverage of all bins instead of between CLB2-064 and individual scaffolds.

### 2.7 Data Availability

All sequence data from this research have been submitted to the NCBI nr and WGS databases under BioProject PRJNA1111131 and BioSamples SAMN41383914 - SAMN41383940. The 16S rRNA reads are available in the SRA database under accessions SRR29010669- SRR29010695. The metagenome reads are available in the SRA database under accessions SRR29054603- SRR29054613. Metagenome assembled genomes are available in the WGS database under accessions SAMN41876582-SAMN41876705 (well A-01); SAMN41876706- SAMN41876780 (well A-02); SAMN41876781- SAMN41876875 (well A-03); SAMN41876876- SAMN41876953 (well A-04); SAMN41877016- SAMN41877096 (well A- 07); SAMN41877097- SAMN41877206 (well B-02); SAMN41877207- SAMN41877317 (well B-03); SAMN41877446- SAMN41877578 (well B-04); SAMN41877579- SAMN41877721 (well B-05); SAMN41877793- SAMN41877969 (well B-06); SAMN41877970- SAMN41878060 (well B-07).

## 3. RESULTS AND DISCUSSION

### 3.1 Aquifer arsenic geochemistry

All samples from Sites A and B fell within circumneutral pH (6.76 – 7.95). Reduction potentials (Eh) ranged from reducing (0 to +200 mV) to highly reducing (-200 to 0 mV), indicative of anoxic conditions (see Table S1 for select geochemistry parameters from 2021 and 2022 groundwater samples).

Dissolved arsenic values in 2021 ranged from 0.016 - 0.035 mg/L in Site A and 0.016 – 0.25 mg/L in Site B. These values were generally stable from 2021 to 2022 (Table S1, Figure 1). The majority of dissolved arsenic is present as As (III), which aligns with the predicted equilibrium of arsenic speciation based on pH and Eh values (Akter et al., 2005).

**Figure 1:**
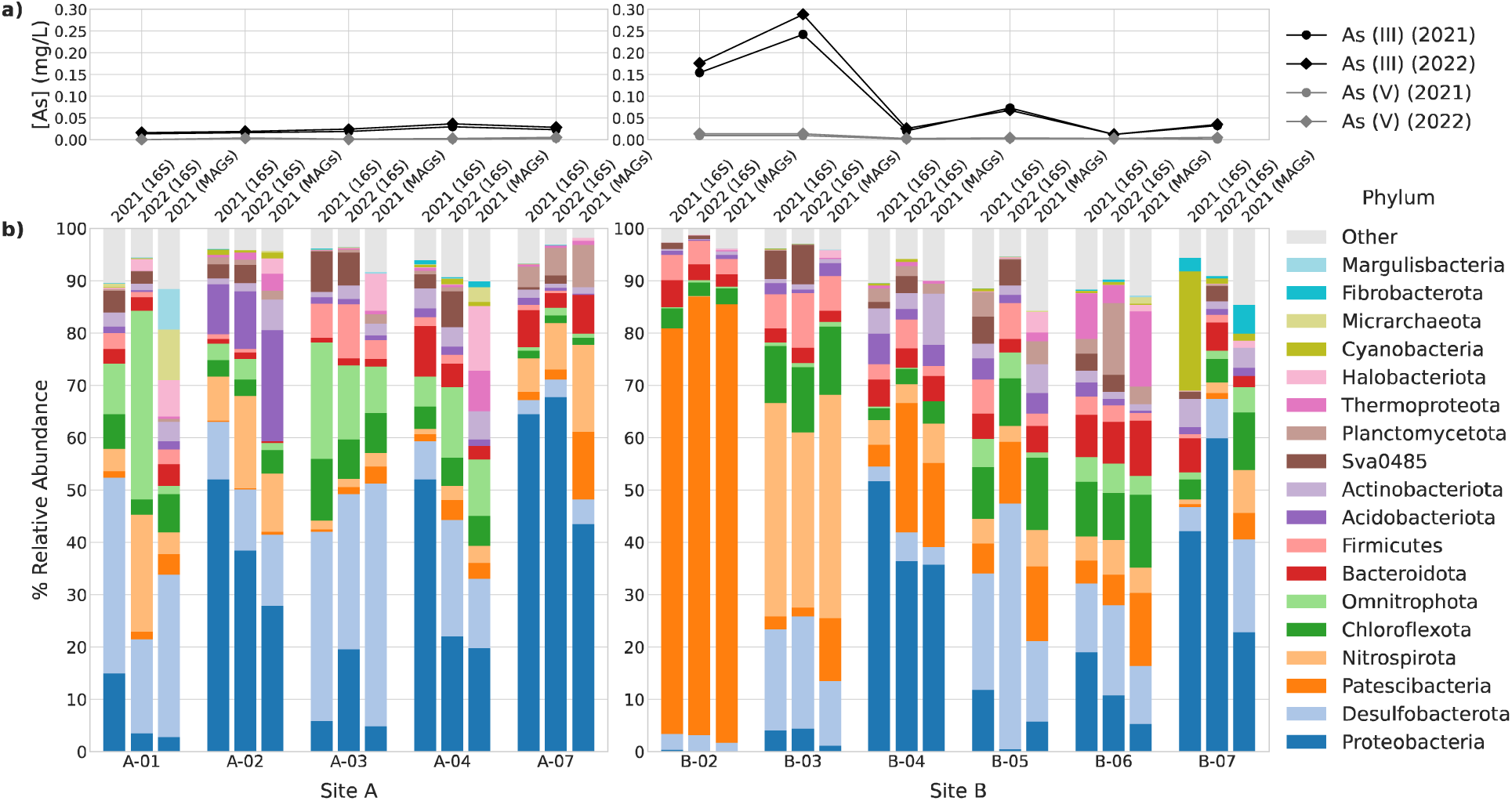
Microbial community composition and arsenic concentrations of groundwater along two aquifer transects. The concentration of dissolved arsenic species (a) for each sample is indicated in mg/L. Barplots (b) indicate phylum-level relative abundances of microbial taxa. Three bars are shown for each sample, representing the 2021 16S rRNA sequencing, 2022 16S rRNA sequencing, and 2021 metagenomic sequencing data in that order. Samples within each site are ordered left to right by the direction of water flow, starting with the well closest to the CSS well pad. Phyla that were present at a relative abundance > 5% in at least one sample in any dataset are shown. Relative abundances were calculated using relative ASV frequencies for 16S rRNA amplicon data and scaffold coverage information for metagenomes. Taxonomy of ASVs were assigned using the SILVA database while MAGs were classified with GTDB-tk. In cases where the nomenclature differed between the databases, the GTDB label is shown. Note that the phylum ‘Sva0485’ (SILVA classification) only appears in the 16S rRNA amplicon datasets. In GTDB this clade is listed under the phylum ‘SZUA-79’, which was not assigned to any of the MAGs; it is likely that the GTDB-tk classifier assigned the related MAGs’ taxonomy under Desulfobacterota, as they are closely related lineages.

### 3.2 Sequencing statistics

A total of 307,306 16S rRNA amplicon reads were processed with QIIME2, resulting in 3,837 ASVs, with a total frequency of 253,383 reads, averaging 23,035 reads per samples (Table 1). Frequencies of individual ASVs across all samples ranged from 2 – 24,217, with a mean frequency of 66.

**Table 1:** 16S rRNA amplicon and metagenomic sequencing statistics by sample. . 16S rRNA amplicon reads were processed in QIIME2 to filter, merge, and group non-chimeric sequences into amplicon sequence variants (ASVs). Assembly of metagenomic reads was performed with SPAdes3. Scaffolds were binned into metagenome-assembled genomes (MAGs) using MetaBAT2, CONCOCT, and MaxBin2, and DAS Tool was used to identify the highest quality set of non-redundant bins among the three binning algorithms. CheckM was used to filter for MAGs with >70% completeness and <10% contamination as the final set of high-quality bins. For detailed information on each MAG see Data File S2.

| Sample | 16S<br>rRNA<br>reads | % Filtered,<br>merged, non-chimeric | # of<br>ASVs | Metagenomic<br>Reads | %<br>Sequences<br>assembled | Scaffolds | Scaffolds<br>binned | High-quality<br>bins |
| --- | --- | --- | --- | --- | --- | --- | --- | --- |
| A-01 | 32,051 | 82.21 | 26,349 | 98,175,715 | 57.26 | 2,254,761 | 23,430 | 124 |
| A-02 | 22,597 | 87.53 | 19,779 | 78,424,298 | 59.17 | 1,980,164 | 14,751 | 75 |
| A-03 | 21,240 | 82.66 | 17,556 | 80,582,181 | 60.94 | 1,747,940 | 16,719 | 95 |
| A-04 | 29,182 | 81.22 | 23,702 | 79,079,594 | 37.83 | 2,817,451 | 19,902 | 78 |
| A-07 | 33,649 | 78.77 | 25,506 | 76,232,243 | 42.54 | 2,404,865 | 12,706 | 81 |
| B-02 | 32,971 | 91.20 | 30,068 | 74,120,634 | 75.90 | 1,231,811 | 18,924 | 110 |
| B-03 | 32,547 | 82.34 | 26,800 | 80,071,928 | 61.69 | 1,838,287 | 22,178 | 111 |
| B-04 | 29,191 | 76.45 | 22,316 | 85,422,107 | 58.94 | 2,254,958 | 18,571 | 133 |
| B-05 | 16,606 | 83.53 | 13,871 | 75,829,097 | 65.89 | 1,678,150 | 25,780 | 143 |
| B-06 | 21,952 | 81.73 | 17,941 | 100,182,557 | 56.48 | 2,677,247 | 29,888 | 178 |
| B-07 | 35,320 | 80.68 | 28,495 | 94,383,591 | 38.46 | 3,412,804 | 17,078 | 91 |
| **Total** | **307,306** | **N/A** | **253,383** | **922,503,945** | **N/A** | **24,298,438** | **219,927** | **1,219** |

We identified 1,219 high-quality MAGs (ranging from 75-178 per sample, average of 107) after assembly and binning of metagenomic sequencing data (Table 1).

### 3.3 Microbial community overview

#### 3.3.1 Community composition

Overall, the taxonomic composition of each well remained relatively stable between 2021 and 2022 samples, as determined by 16S rRNA ASV frequencies (Figure 1). The 2021 metagenomic composition data, determined by average scaffold coverage of MAGs, was also consistent with the 2021 ASV frequencies (Figure 1). Well B-07 was a slight exception, with a high abundance of Cyanobacteria ASVs (classified under f. Nostocaceae) detected in 2021 (see Figure 1). This unexpected result could be explained by contamination by plant matter, where plastid DNA was amplified during 16S rRNA amplicon library preparation.

Community composition among samples collected in Site A were generally similar at the phylum-level, which was expected based on the stability of geochemical conditions across the transect following migration of the arsenic plume downgradient (Table S1). Arsenic concentrations were relatively low in all wells but slightly above background concentrations (0.022 mg/L) in wells A-02, A-04, and A-07, as the site has not experienced recent steaming. Well A-07 was most recently impacted by the plume (2013), and had a higher proportion of Proteobacteria (68% in 2022 vs. 4-38%) and lower proportion of Chloroflexota/Omnitrophota (< 2% in 2022 vs. 3-36%) compared to other Site A wells (Figure 1). It is possible that the community hadn’t stabilized at well A-07 at the times of sampling, or that the slightly higher redox potential (Table S1) supports different microbial guilds at this location (discussed below).

In contrast, Site B wells had high variability in both arsenic concentration and the dominant phyla present (Figure 1). B-02 and B-03 were experiencing elevated arsenic concentrations (0.15-0.25 mg/L) at the time of sampling from the most recent steam cycle, which held stable across 2021 and 2022 due to the slow flow rate (< 5 m/year) of the aquifer. Well B-02 contained a high relative proportion of Patescibacteria (∼80%) that was not observed in any other sample, mainly from a single population within the Class Paceibacteria. GTDB-tk classified the Paceibacteria MAG (CLB2-064) under family/order UBA6257; other members of this clade have primarily been found in groundwater environments (Zhao et al., 2022). The Patescibacteria (formerly Candidate Phyla Radiation) is a recently described phylum, whose members are characterized by small cell sizes and highly reduced genomes, with many metabolic pathways incomplete or absent (Brown et al., 2015; Fujii et al., 2022; Tian et al., 2020). They are believed to be obligate symbionts (likely epibiotic parasites) of other bacteria and/or archaea (Kuroda et al., 2022).

Well B-03 also had a distinct community composition compared to all other samples. Well B-03 was dominated by 2 taxa within the phylum Nitrospirota, with MAG relative abundances of 15.43% and 24.54% respectively. Both MAGs (CLB3-019 and CLB3-097) were classified under order Thermodesulfovibrionales (c. Thermodesulfovibrionia), which contain obligately anaerobic, thermophilic sulfate reducers (Umezawa et al., 2021). Both MAGs were classified to unnamed families. Wells B-04, B-05, and B-06 had similar microbial compositions across both years. As with Site A, higher concentrations of arsenic previously occurred in these wells (*i.e.,* from 2005-2010), and the communities appear to have stabilized following a decrease in arsenic concentration. Notably, the community composition in Well B-07 shows similar phylum-level groups but differing abundances compared to Wells B-04, 05, and 06. Well B-07 is much farther downgradient from the CSS well pad than the other wells and has not yet experienced elevated arsenic concentrations. It is therefore indicative of the aquifer communities prior to steaming and the resultant heat and arsenic exposures. While wells B-04, B-05, and B-06 appear to be returning to this background community composition, there remains substantial disruption to the original groundwater communities.

#### 3.3.2 Diversity

All diversity metrics were calculated from 16S rRNA amplicon sequencing data, with both 2021 and 2022 samples included.

Overall, there were no significant differences in α-diversity between sites (sampled within the same year) and between samples of the same aquifer in different years (Figure S1). Well B-02 is consistently an outlier, being the lowest in α-diversity, due to the single Patescibacteria ASV comprising the majority of the community (Figure 1, Figure S1). Well B-03’s α- diversity is considerably higher compared to B-02 (comparable to other samples in Site B), despite having the highest total arsenic concentration, indicating that arsenic concentration itself is not deterministic of a severe reduction in community richness/evenness.

A correspondence analysis was performed to compare the *X*^2^ distances between the samples and features (ASVs) (Figure S2). Correspondence analysis supports the community composition data (Figure 1) indicating that the A-07, B-02, and B-03 community compositions are highly divergent from each other, represented by the distance between the samples on the ordination axes. Furthermore, the ASVs that were highly abundant in wells A-07, B-02, and B-03 also diverged from each other and clustered near their respective sample on the biplot (Figure S2).

β-diversity was evaluated with principal coordinates analysis (PCoA) using Bray-Curtis and unweighted/weighted unifrac distance matrices (Figure S3). Unlike α-diversity, all β- diversity ordinations showed a significant difference between sites (ANOSIM, p<0.05), indicating that the specific microbial populations present at Sites A and B differ. There were no significant differences between years.

To evaluate the relationship between community composition and environmental variation across samples, we projected geochemical factors onto PCoA ordination plots as loading vectors (Figure 2, Figure S3). The unweighted unifrac plot (Figure 2) includes a subset of geochemical variables (twelve total, listed in Table S1), which were selected based on the following criteria: 1) variables that were measured in both 2021 and 2022, 2) values were above the detection limit in all wells, and 3) the variable was either predicted to influence arsenic biogeochemical cycling (*e.g.*, Fe, SO_4_) or major microbial metabolisms (*e.g.*, available C/N species). Significant variables were determined by environmental fitting (see Methods) of values to PCoA ordinations.

**Figure 2:**
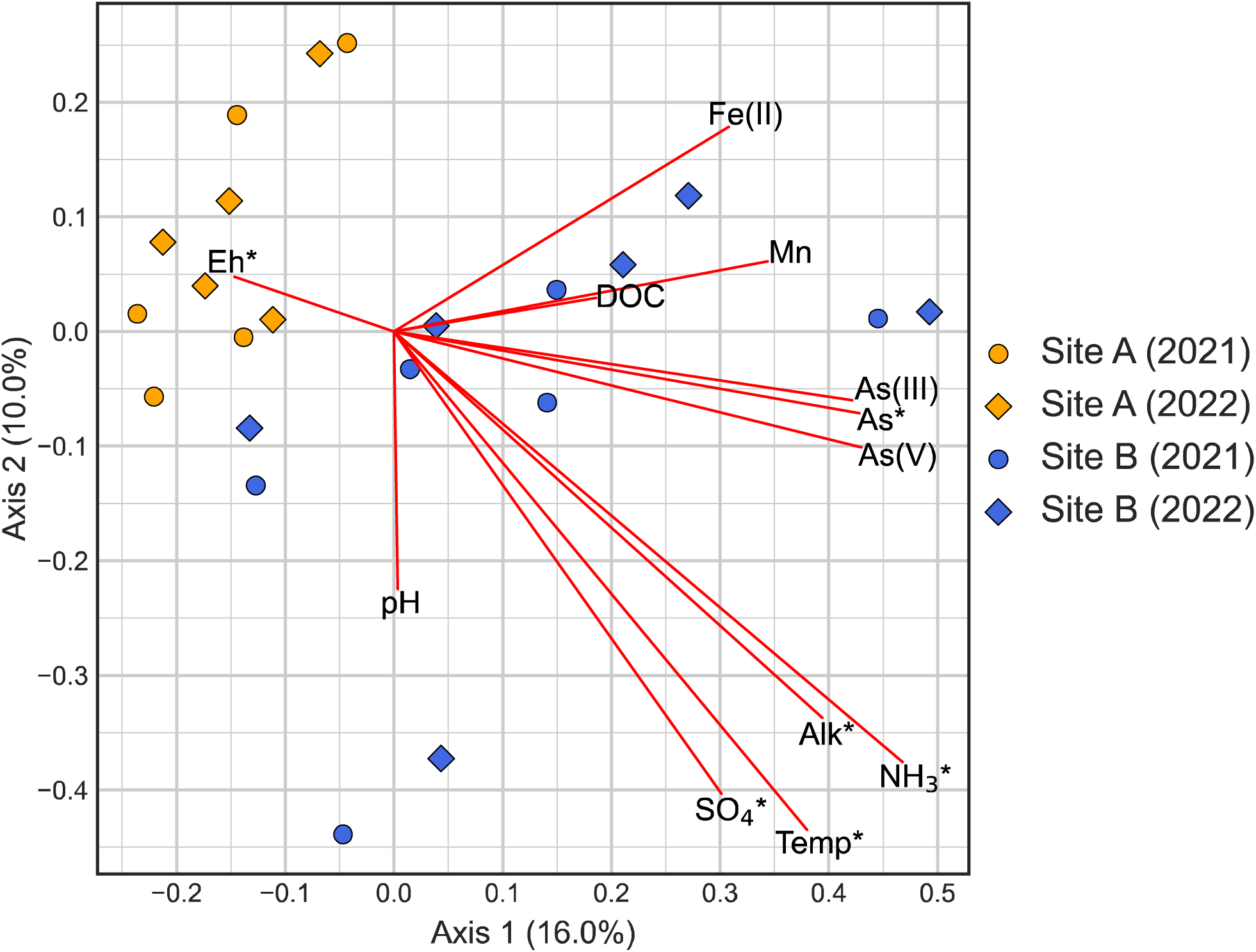
Principal Coordinates Analysis (PCoA) biplot of unweighted unifrac dissimilarity. Samples are indicated by points, with colors representing sites and shape representing year. Axis labels indicate the proportion of variation explained. Vectors (red) indicate the projection of selected environmental factors onto the ordination, and are scaled to the maximum/minimum values of each axis. Significant variables (p<0.05, see Table S1) are labelled with an asterisk. Abbreviation key: Temp = Temperature, Alk = Alkalinity (in CaCO_3_), As = Total Dissolved Arsenic, DOC = Dissolved Organic Carbon. The unweighted unifrac distance matrix was chosen over other PCoA ordinations, as the most environmental variables were found to significantly correlate with the ordination (Table S1). See Figure S3 for ordinations that display all measured geochemical variables.

Six geochemical factors significantly correlated (p<0.05) with at least one ordination: Eh, temperature, total arsenic, ammonia, alkalinity, and sulfate (Table S1). Several trace metals and other ions also showed significant correlations with β-diversity (Data File S2), but are not displayed in Figure 2, as they did not meet the 3^rd^ criterion listed above. It is difficult to conclude whether these variables exert any direct influence on community diversity, as it is not clear whether their correlation with community dissimilarity is due to covariance with another environmental variable. Figure S3 shows all ordination plots with all variables projected.

Samples primarily separate based on redox potential (higher in Site A) and sulfate/ammonia/alkalinity (higher in Site B) (Figure 2). Temperature also drives separation of Sites A and B (generally higher in Site B, likely due to more recent steaming and slow water flow) (Figure 2), but also accounts for much of the variability between samples within Site B (6.9-32.6 °C in Site B vs. 10.5-15.3°C in Site A) (Table S1). Total arsenic concentrations play a comparatively minor role in predicting community diversity between sites and samples (p=0.041-0.144, Table S1), which is consistent with the fact that arsenic resistance/metabolism genes are widely distributed among bacteria and archaea (Crognale et al., 2017; Zhu et al., 2017). Therefore, while localized instances of elevated arsenic correlate with short-term changes in community composition (see Figure 1), a key selective force impacting aquifer communities appears to be the increase in water temperature in the wells closest to the CSS well pad, as a result of heat conduction from steam injection. Additionally, the succession of long-term, stable communities following the dispersal/migration of thermal and arsenic plumes is primarily dependent on natural aquifer redox conditions and nutrient availability.

### 3.4 Distribution and abundance of arsenic-related genes

#### 3.4.1 Direct arsenic-cycling genes

To predict the microbial capacity for direct arsenic cycling in each aquifer, annotation of arsenic-related genes was performed on all MAGs. Efflux pumps (*ars*B/*acr*3), intracellular arsenic reductase (*ars*C), and arsenic methyltransferase (*ars*M) were common in nearly all phyla (Figure 3). Exceptions to this were the Patescibacteria and members of the archaeal DPANN superphylum (Micrarchaeota and Nanoarchaeota), all lineages characterized by small, reduced genomes. Dissimilatory As(V) reduction, indicated by *arr*A, was uncommon and restricted to members of the Nitrospirota, Desulfobacterota, and Proteobacteria phyla (Figure 3). As(III) oxidation genes (*aio*AB, *arx*A) were the most rare: only 7 MAGs encoded these genes, all of which were from members of the Proteobacteria (Figure 3).

**Figure 3:**
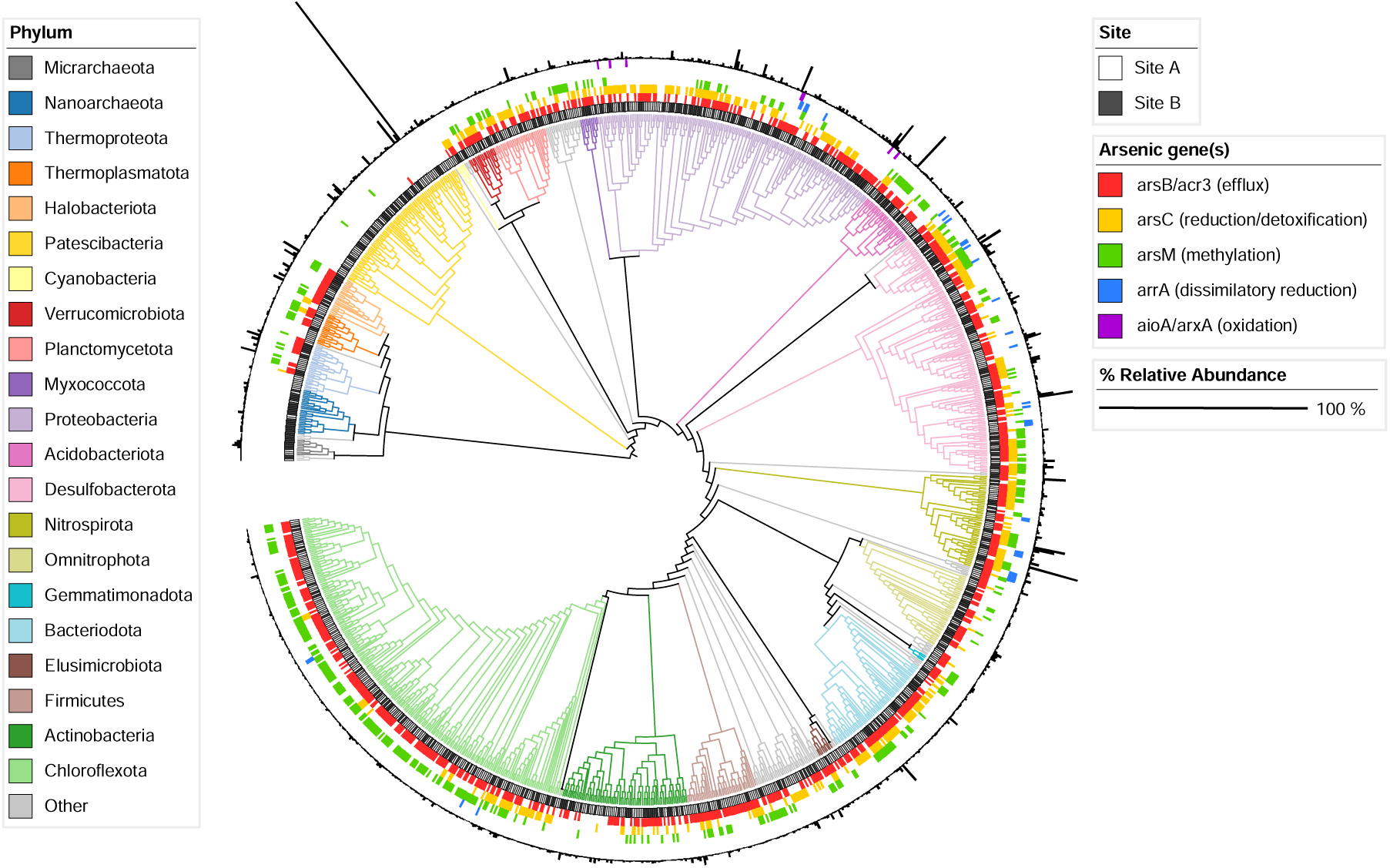
Phylogenetic distribution of arsenic-related genes across all MAGs. Tree branches are colored by phylum in clockwise order starting at the Archaea (∼10 o’clock). The ‘Other’ category represents phyla with < 3 representative MAGs in the total dataset. The innermost ring designates which site the MAG originated from. Colored rings (ordered by innermost to outermost on the legend) represent presence/absence of arsenic-related genes, where at least one hit for any gene within each category was used to determine presence. The outermost ring is scaled to the % relative abundance of each MAG within its sample.

The presence/absence of arsenic-cycling genes across MAGs (Figure 3) may not reflect the proportion of community members possessing these genes, due to variation in species abundance. To compare arsenic gene abundance across samples, positive hits for each gene (presence of at least one copy) were aggregated for MAGs within each sample and normalized to the relative abundance of MAGs (Figure 4). Across all samples, arsenic resistance and methylation genes were more abundant compared to metabolic arsenic reductase/oxidases (Figure 4). Efflux pumps within MAGs were mostly encoded by *acr*3, which has been previously found to be more widespread than *ars*B (Zhu et al., 2017).

**Figure 4:**
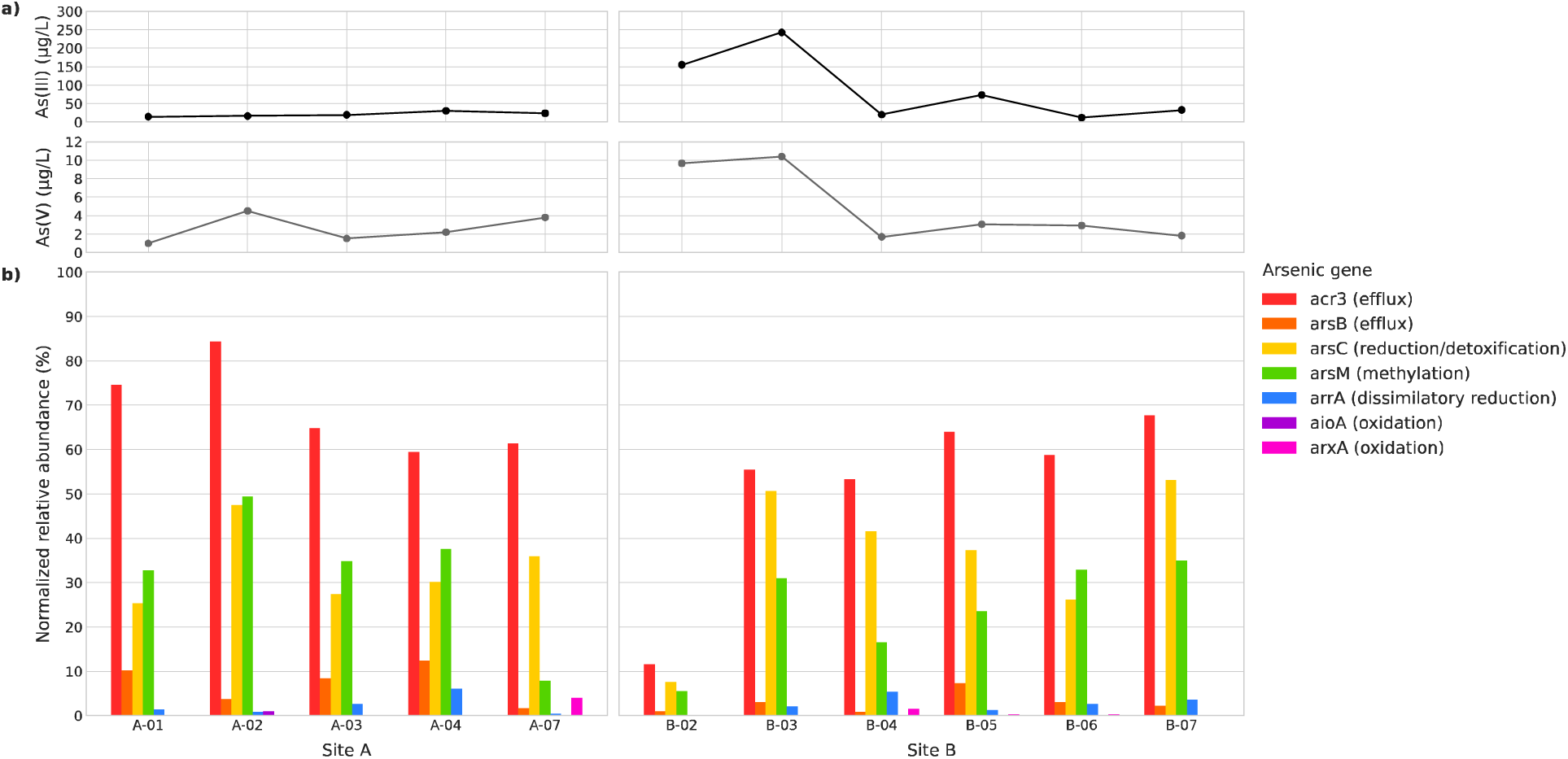
Predicted abundance of arsenic-related genes across wells. The concentrations of dissolved As(III) and As(V) are shown in a), and abundances of arsenic-related genes in corresponding samples are shown in b). Normalized abundances of genes were calculated by taking the sum of the relative abundances of MAGs that were positive for one or more hits of that gene. Normalized abundances therefore also represent the percentage of organisms present in each sample predicted to be capable of that function.

Arsenic concentrations within wells did not correlate with an increase in abundance of arsenic-cycling genes. Well B-02 had the second highest arsenic concentration but the lowest overall abundance of all As-cycling-associated genes (Figure 4). This trend is attributable to the dominant population of Paceibacteria (discussed further in section 3.4.3). B-03 shows similar gene abundance profiles compared to the rest of the aquifer despite having the highest arsenic concentration (Figure 4). These observations align with our whole-community analyses and suggest that arsenic concentration has a minor bearing on the distribution of arsenic-specific pathways.

In this instance, the disconnect between arsenic concentration and the distribution of arsenic cycling genes can be attributed to the fact that arsenic resistance/metabolism genes are widely distributed among bacteria and archaea (Crognale et al., 2017; Zhu et al., 2017). Additionally, the *aio*, and *acr*3, and *ars* operons all have associated regulatory genes that activate or repress transcription based on As(III)/As(V) concentrations (Romero et al., 2022; Shi et al., 2020; Suzuki et al., 1998). We suspect that if the microbial community within B-03 is more resistant to arsenic toxicity, this resistance may be tied to the upregulation of *acr/ars* genes. It is also possible that As(III) oxidizers are expressing *aio*AB/*arx*A at higher rates relative to the oxidizers’ abundance. Metatranscriptomic and/or metaproteomic data would be required to confirm changes in rates of arsenic cycling activity within the aquifers.

The *arsM* gene was the third most abundant arsenic-related gene detected overall (Figure 4). Biomethylation is thought to contribute to the cycling of arsenic between aquifers and surface environments, due to methylarsines being more volatile than inorganic species and more susceptible to diffusion (Maguffin et al., 2015). It is possible that microbial arsenic methylation via ArsM provides an additional mechanism of arsenic removal from the system, from volatilization of organic arsenicals (Crognale et al., 2017). However, it is unlikely to be a major factor impacting arsenic cycling at Sites A and B. If organic arsenicals were being produced in this system, we would not expect them to readily escape via volatilization given the physical barriers inherent in the aquifer (*e.g.,* cut off from the atmosphere and slow groundwater flow). We did not detect measurable concentrations of mono- or dimethylarsenic acid in any sample, suggesting ArsM is not active despite its widespread distribution.

The rarity of As(III) oxidizers can be explained by the restricted niche space these guilds encounter in this system. While *aio*/*arx* genes are widely distributed across the bacterial domain, all known As(III) oxidizers are either aerobes (*e.g.*, *Rhizobium, Thiobacillus, Chloroflexus, Alcaligenes* spp.) (Hamamura et al., 2023; Inskeep et al., 2007; Rhine et al., 2007) or anaerobes that couple As(III) oxidation to denitrification (*e.g.*, *Alkalilimnicola ehrlichii* MLHE-1, *Sinorhizobium* spp.) (Oremland et al., 2002; Rhine et al., 2007; Zargar et al., 2010). Redox potential (Eh) measurements indicate that all wells are likely anoxic (Table S1) in addition to nitrate being below detection limit everywhere except A-07 (0.51 mg/L). Notably, A-07 showed the highest prevalence of *arx*A (encoded by a novel *Gallionella* MAG, CLA7-040, at 4.01% abundance), which supports our hypothesis that low availability of compatible electron acceptors limits the distribution of As(III) oxidizers.

Thermodynamic calculations for As(III) oxidation coupled to the reduction of other electron acceptors available (e.g., Fe(III) or SO_4_^2-^) indicated these metabolic processes are not thermodynamically favourable (*i.e.*, ΔG > 0) at As(III) concentrations and temperatures representative of this study site (*i.e.*, 0.1 to 100 µM As(III) and 0 to 200 °C) (Basu et al., 2010). Although a recent study showed that As(III) oxidation coupled to Fe(III) reduction does occur abiotically under anoxic conditions (Fang et al., 2023), there is no reported physiological evidence of microbial As(III) oxidation coupled to the reduction of iron-bearing minerals, suggesting it is unlikely to be occurring at this petroleum extraction site.

In summary, the overall trend of microbial activity with respect to direct arsenic cycling in both aquifers heavily favors intracellular reduction of As(V) and transport of As(III) out of the cell, which increases the relative concentration of the more toxic/mobile As(III) species. Although arsenic methylation pathways were widely detected, there is no geochemical evidence they are active in this system. Finally, As(III) oxidation pathways that could counteract the mobilization of arsenic were rare, likely due to a requirement for oxygen or nitrate in a system lacking both.

#### 3.4.2 Iron- and sulfur-cycling genes

The biogeochemical cycling of arsenic is directly connected to iron and sulphur redox cycling in the environment. Due to these interactions, we considered how biological iron and sulphur redox cycling could control the fate of arsenic in a system dominated by arsenic detoxification strategies that favour the reduction of As(V) to As(III). Annotated Fe/S redox genes were aggregated and normalized to the abundance of MAGs within each sample, reflecting the proportion of organisms within each community that encode these genes (Figure 5).

**Figure 5:**
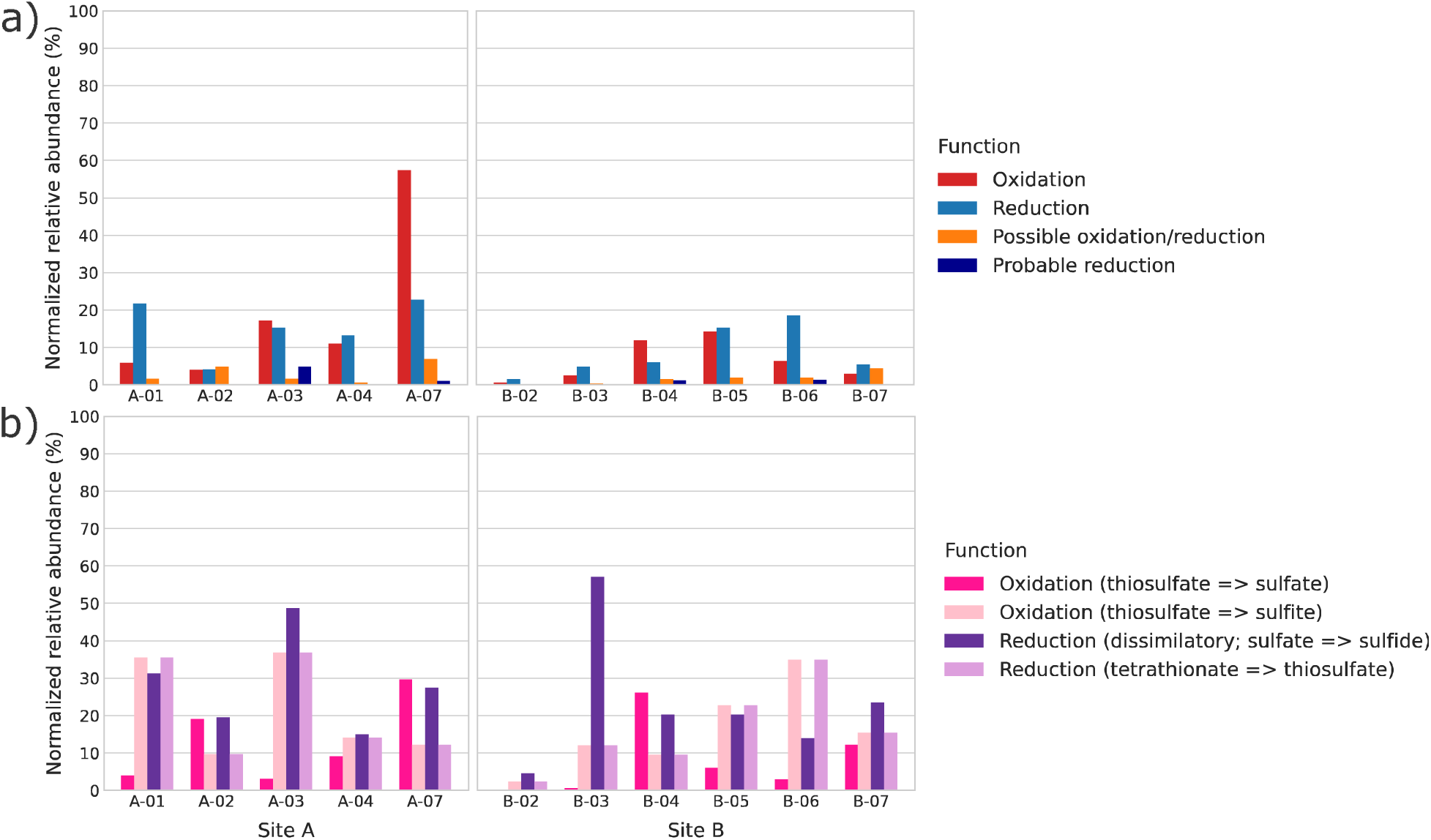
Predicted abundance of a) iron and b) sulfur cycling genes across wells. Genes are categorized by function (oxidation and/or reduction) for both elements. Normalized abundances for functions were calculated by taking the sum of the relative abundances of MAGs that were positive for one or more gene within the function (and therefore represent the percentage of organisms present in each sample predicted to be capable of that function). Gene families were identified and categorized using FeGenie (iron) and DRAM (sulfur). Iron oxidation genes include *cyc2, fox* genes, and sulfocyanin; reduction genes include *omc* and *dfe* genes. Other hypothesized (but not verified) genes for iron cycling include *mto* (possible oxidation/reduction) and *mtr* (probable reduction) genes, which are indicated separately (orange and purple, respectively). A full list of iron-related genes used in this analysis can be found in Garber et al., 2020. Sulfur oxidation genes are divided into the SOX pathway (thiosulfate to sulfate) and other pathways that produce sulfite as an intermediate. Dissimilatory reduction of sulfate is identified *dsr* and *apr* genes, while reduction of tetrathionate to thiosulfate is identified by the *ttr* gene.

Aerobic Fe(II) and sulfur oxidation is unlikely to be occurring within most wells in Sites A & B, even if genes for these metabolism were detected in the metagenomes (Figure 5). Given redox potentials are below 200 mV (Table S1), it is improbable that enough oxygen is present to support the oxidation of ferrous iron and reduced inorganic sulfur compounds. Simultaneous oxidation of elemental sulfur and As(III) has been demonstrated in some species, but requires aerobic, oxidizing conditions such as those present in alkaline lake waters (Fisher et al., 2008), which is therefore not applicable to groundwater arsenic cycling (pH 6.76-7.95, Table S1). The maximum Fe(III) concentration measured was 0.41 mg/L (B-02), and the maximum H_2_S concentrations measured was 0.15 mg/L (B-04), which do not correlate with abundances of iron oxidizing and sulfate reducing genes, but we acknowledge that groundwater concentrations are not indicative of Fe(III)/H_2_S production due to their poor solubility (Lux, 1963; National Center for Biotechnology Information, 2024).

The community within A-07 may be an exception, as over 50% of organisms within this sample were predicted to have iron oxidation genes (Figure 5b). Most of these genes were represented by novel MAGs belonging to the genus *Gallionella*, which are known microaerophilic neutrophilic Fe(II)-oxidizing bacteria. The abundance of *Gallionella* in A-07 is the main factor driving its community dissimilarity compared to the other wells (Figure S2), as this group is largely absent from other samples. It is possible that small amounts oxygen infiltration could support microaerophilic Fe(II) oxidation by *Gallionella* spp. in A-07, or that they are coupling this metabolism with NO_4_^-^ reduction, which has been observed in members of the family Gallionellaceae (Huang et al., 2021, 2022). If this is the case, we would expect Fe(II) oxidizers to have low activity due to the same niche restrictions highlighted for As(III) oxidizers.

Reductive iron and sulfur metabolisms are more favored in anoxic reducing environments, such as the deep aquifers in this study. The presence of Fe(III) reduction capacity (present in ∼20% of organisms in some wells) presents a possible mechanism for arsenic mobilization, as reductive dissolution of Fe(III)-bearing minerals can lead to de-sorption of arsenic from sediments (Smedley & Kinniburgh, 2002). The production of biogenic sulfides from sulfate-reducing bacteria (SRBs) can have a variety of indirect effects on the arsenic cycle. Under reducing conditions and low dissolved iron (such as this study’s system), sulfide may form stable thioarsenite complexes with As(III), which increases its mobility (Bostick et al., 2005; Wilkin et al., 2003). Other reactions may simultaneously occur that form arsenic sulfide minerals (AsS, As_2_S_3_) that limit arsenic’s mobility through precipitation (Fisher et al., 2008; Newman et al., 1997; Wilkin et al., 2003). The highest arsenic well, B-03, also has the highest abundance of SRBs (Figure 5b), which is attributed to the Thermodesulfovibrionia MAGs in that community, the most prominent populations being CLB3-097 (24.5% relative abundance) and CLB3-019 (15.4% relative abundance) (Data File S2). Given the comparatively low abundance of sulfur oxidizers relative to SRBs (Figure 5b) and the low groundwater Eh (Table S1) in B-03, we predict that biogenic production of sulfide due to SRB may be one of the main pathways limiting arsenic’s mobility in the system.

#### 3.4.3 Potential host-symbiont interactions and arsenic resistance mechanisms of Paceibacteria CLB2-064

The high abundance of CLB2-064 within well B-02 was highly unexpected for two reasons. First, the genome of CLB2-064 is consistent with other members of the Paceibacteria, with pathways for glycolysis, the TCA cycle, and electron transport missing or incomplete (evaluated using DRAM, Data File S2), suggesting an obligately symbiotic or epibiotic lifestyle. Secondly, this population was found to be dominant in a well with higher arsenic concentrations (0.17 mg/L in 2021), despite lacking any known arsenic resistance genes.

To identify potential host-symbiont interactions involving CLB2-064, proportionality (or co-occurrence) analysis was performed on CLB2-064 MAG coverage compared to the coverage of B-02 scaffolds across all samples. Scaffolds showing proportional abundance to CLB2-064 abundance could represent potential hosts for this species, with a shared abundance pattern but not shared absolute abundance levels.

All scaffolds >10kb that correlated with CLB2-064 MAG coverage belonged to existing MAGs (Data File S2). A subsequent proportionality analysis was performed between the relative abundance of CLB2-064 and all other MAGs in B-02, with only one MAG identified to be proportional to CLB2-064 at the full genome rather than scaffold level (CLB2-037, Data File S2). Proportionally abundant scaffolds and MAGs were taxonomically diverse, and present at a maximum abundance of 1.349%. It is unlikely that any of these MAGs form a single host population for CLB2-064 given these abundances; even if multiple parasitic cells attach to a single host cell (as shown in Kuroda et al., 2022), the ratio of symbiont to host would be greater than 60:1. We initially considered the possibility that CLB2-064 dominance in 2021 samples may have been due to an event just prior to sampling that killed its host population, and the CLB2-064 population had not yet fallen in response. However, the 2022 16S rRNA amplicon data showed that the community composition remained stable over at least one year. Given these results, we hypothesize that Paceibacteria CLB2-064 can either engage in symbiosis with low host specificity and are able to parasitize multiple other species present in the aquifer, or that they parasitize a eukaryotic host that was not captured in this analysis. Patescibacteria have been identified as intracellular symbionts in ciliates, a dynamic that may be occurring in this system as well (Gong et al., 2014). Future research could use microscopy (specifically cryo-TEM as used in (He et al., 2021), which is able to image Patescibacteria mechanisms of attachment to host cells) and/or culture-based methods to elucidate the role of these understudied organisms within the aquifer community.

The ability of CLB2-064 to survive high concentrations of arsenic despite not encoding *ars* or *acr* genes may be a by-product of the characteristically reduced genomes in this clade, rather than the evolution of a completely different mechanism of arsenic resistance. The necessity for intracellular As(V) reductases and membrane As(III) efflux pumps is due to the fact that arsenic must enter the cell in order to be cytotoxic. As(III) has been shown to enter through aquaglyceroporins (*e.g.*, glycerol facilitator GlpF), while As(V) is very similar in structure to phosphate and enters through phosphate uptake systems (*e.g.,* Pit, Pst) (Crognale et al., 2017; Kruger et al., 2013). Gene annotation data for the CLB2-064 MAG showed that both glycerol and phosphate transporter genes were completely absent. We hypothesize that arsenic resistance in CLB2-064 is a byproduct of their lack of membrane transporters for As(III) and As(V) to exploit to enter their cells. Alternatively, if CLB2-064 is an intracellular parasite to a ciliate, it may rely on the host’s arsenic resistance mechanisms, which also eliminates the need to encode them in their own genome

### 3.5 Opportunities for remediation

#### 3.5.1 Arsenic oxidation

As discussed in 3.4.1, As(III) oxidizers are rare in this study system, most likely due to the absence of electron acceptors (O_2_, NO_4_^-^) for As(III) oxidation in deep, anoxic groundwater environments (low redox potential). Leveraging As(III) oxidizing guilds for bioremediation would likely require supplementing the aquifers with NO_4_^-^ and potentially introducing large exogenous populations of As(III) oxidizers into the aquifers. These methods are unlikely to be efficient at counteracting the mobilization of arsenic in this system because the widespread *arsC* detoxification pathways would reduce As(V) back to As(III). Moreover, the capacity of exogenous As(III) oxidizers to survive the steep physicochemical gradients (*e.g.*, temperature fluxes due to in-situ bitumen recovery processes) encountered in the aquifer would need to be considered as part of whether this is feasible.

#### 3.5.2 Formation of iron minerals

One Fe(II)-oxidizing *Gallionella* MAG (CLA7-040) was the most abundant As(III) oxidizer. As(V) and Fe(III) can co-precipitate as scorodite under oxidizing conditions, so stimulating both metabolisms (potentially even in the same organism) could support arsenic removal. We expect such a strategy would find limited purchase because scorodite is relatively soluble in circumneutral pH conditions (Langmuir et al., 2006). Although scorodite formation could mitigate extremely high arsenic levels, the equilibrium of dissolved/solid phase arsenic would still be higher than regulatory standards for groundwater.

Adsorption of arsenic onto Fe(III) minerals (mainly iron (oxyhydr)oxides) is a more effective mechanism of controlling arsenic mobility in neutral pH groundwater. Increasing the rate of Fe(III) mineral formation through stimulating microbial Fe(II) oxidation may be possible through the addition of NO_4_^-^. Additional Fe(II) would also need to be added, as natural concentrations (0.28-2.5 mg/L) within the aquifers are low in the context of bioavailability as an electron donor (Table S1). Redox conditions within these aquifers do not currently favour biological oxidation of Fe(II) anywhere except A-07. For all other locations, the capacity to amend sufficient oxidized species to raise the redox potential and stimulate iron oxidation will likely be a key limitation to consider with such a strategy.

#### 3.5.3 Formation of sulfide minerals

As discussed in 3.4.2, sulfide may increase arsenic mobility by forming thioarsenite complexes with As(III) (Bostick et al., 2005; Wilkin et al., 2003). Arsenic may also precipitate with sulfides as arsenic sulfide minerals (AsS, As_2_S_3_), but similar to scorodite, the equilibrium of arsenic with these minerals at pH 7 is not favored towards the solid phase (Saunders et al., 2018); the stimulation of SRBs alone is therefore not reliable for large-scale remediation of thermally mobilized arsenic plumes.

If SRB activity is stimulated with the combined addition of Fe(II), the formation of pyrite is more favorable – leading to subsequent sorption of As, and incorporation of As(III) into growing pyrite crystals (arsenopyrite) (M.-K. Lee et al., 2019; Saunders et al., 2018). A field-scale remediation experiment demonstrated this principle, where researchers amended arsenic-contaminated groundwater with sulfate, ferrous iron, along with organic C/N/P sources (M.-K. Lee et al., 2019; Saunders et al., 2018). The treatment was effective at sequestering arsenic via the formation of biogenic pyrite, reducing dissolved concentrations from 0.5 mg/L to <0.05 mg/L.

Sulfate concentrations within Sites A & B range from 22-59 mg/L (Table S1), making them comparable to initial concentrations in the field remediation experiment described above. To replicate their results, additional sulfate would need to be added to the aquifer (up to 300 mg/L), as well as ferrous iron (up to 100 mg/L), and labile C/N/P sources to support the growth and metabolism of native SRBs that are widely distributed in this system.

The Thermodesulfovibrionia MAGs identified in this study (e.g., Thermodesulfovibrionia CLB3-097 and CLB3-019) may be particularly useful for remediating aquifers impacted by steam extraction of bitumen because they are able to tolerate high arsenic concentrations and heat (but appear to be abundant once temperatures have lowered back down to ∼25°C as well). Thermophilic SRBs that are active at high temperatures but tolerant of lower temperatures may be better suited to counteracting arsenic plumes during active steaming and shortly after before the heat has dissipated. Mesophilic SRBs that can persist through cycles of steam injection and become active once temperatures lower would be more suited for long term remediation of arsenic-impacted aquifers where steam injections have ceased but arsenic levels remain high.

In summary, the biogeochemistry of groundwater across Sites A & B favors biological sulfate reduction as a major redox reaction driving indirect arsenic cycling. The high arsenic well B-03 appears to support a niche containing arsenic-tolerant, thermotolerant SRBs that show potential for remediation of thermally mobilized arsenic, although biostimulation via the addition of sulfate, iron, and nutrients would be required.

## 4. CONCLUSIONS

In this study, we profiled the taxonomic composition and diversity of aquifer communities experiencing recent elevated arsenic concentrations with respect to background due to *in-situ* thermal bitumen extraction. Metagenomic sequencing was also used to predict the functional capacity of microbes to contribute directly to arsenic cycling and indirectly control the fate of arsenic through iron and sulphur cycling. The main goal was to identify organisms and/or consortia that could mitigate arsenic’s mobility under the geochemical constraints of our study site.

Community composition was highly variable between higher-arsenic and lower-arsenic wells, although the dominant taxa within higher-arsenic wells were also different from each other within and between transects. Environmental factors such as Eh, temperature, alkalinity, [SO_4_^2-^], and [NH_3_] were found to correlate more strongly with community diversity than arsenic concentrations, indicating that groundwater communities stabilize differently after short-term arsenic exposure based on the native geochemical conditions of the aquifer.

Genes for arsenic resistance (*ars*B, *acr*3, *ars*C) and methylation (*ars*M) were widespread in all samples, regardless of current arsenic concentrations. Microbial Communities predominantly contain organisms predicted to reduce intracellular As(V) and produce As(III), which could release aqueous As(III) complexes back into solution. In contrast, arsenic metabolic genes (*arr*A, *aio*A, *arx*A) were rare. Arsenic oxidation is not likely to be supported in these environments due to limited terminal electron acceptors suitable for these metabolic reactions.

Iron and sulfur-cycling organisms were also identified that may indirectly contribute to arsenic attenuation. In more oxidizing environments (such as A-07), iron oxidizing *Gallionella* spp. could immobilize dissolved arsenic by favouring the precipitation of ferric (oxy)hydroxide minerals that strongly sorb arsenic. In reducing environments, sulfate-reducing bacteria, such as the novel Thermodesulfovibriales MAGs within the highest-arsenic well, can induce sulfide mineral formation, which could remove arsenic by co-precipitation (pyrite/arsenopyrite). Given the reducing conditions of most wells in the studied aquifer system, we predict that stimulating SRB activity with the addition of sulfate, along with ferrous iron, carbon, nitrogen, and phosphorus, is a potential strategy for bioremediation of thermally mobilized arsenic.

An unexpected finding of a highly abundant Paceibacteria population in one high arsenic well could not be fully explained, as there was not a clear host population capable of supporting a symbiont present at >80% relative abundance. This population can persist at high abundances despite not encoding arsenic resistance genes. If they are endosymbionts of an unidentified eukaryote, they may benefit from arsenic resistance mechanisms of their host. Otherwise, the fact that their genome lacks genes for the membrane transporters that bring arsenic into the cell may constitute an alternative arsenic resistance strategy.

In summary, aquifer communities experiencing past and present short-term elevated arsenic consist of a wide range of organisms, due to arsenic resistance mechanisms being widespread among microbes. The high taxonomic and metabolic diversity observed across aquifer transects that differ in arsenic concentrations as well as other environmental factors present an opportunity to further explore microorganisms that directly and indirectly influence arsenic mobility in groundwater.

## ACKNOWLEDGEMENTS

We would like to thank our collaborators at Imperial Oil Resources Ltd., including Asfaw Bekele and Logan Swaren for their insight, fieldwork support, and interest in this project. This work was supported by Imperial’s University Research Award. MC was supported by an NSERC CGS-D graduate scholarship. DSG was supported by a Banting Postdoctoral Fellowship from NSERC. LAH was supported by a Tier II Canada Research Chair.

