## Supplemental materials for "Aquifer microbial communities are highly variable in response to thermal arsenic mobilization"

### SUPPLEMENTARY DATA

Table of Contents

Figures S1-3	p. 2-4

Table S1	p. 5-6

Data files 1 and 2 info	p. 7

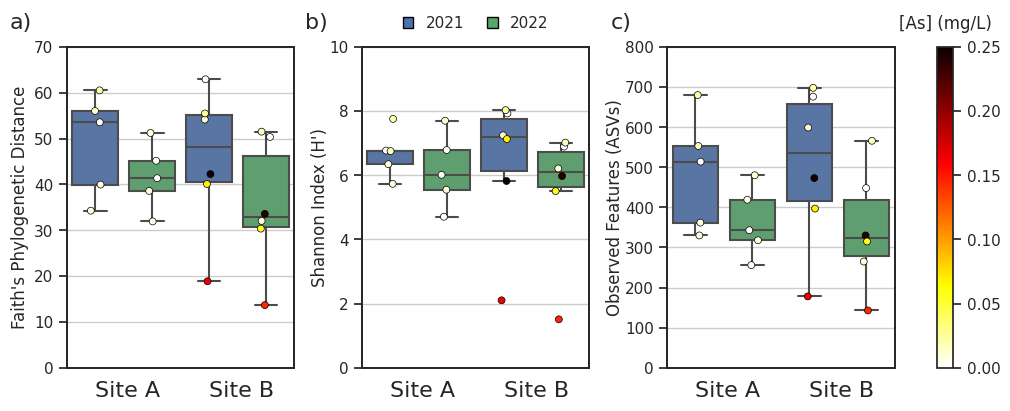

**Figure S1: Alpha diversity of groundwater samples from two aquifer transects.** Measures of a) Faith’s phylogenetic distance, b) Shannon diversity index, and c) number of observed ASVs are summarized for samples from each aquifer (Site A and Site B), taken from the same set of wells in 2021 and 2022. Data was rarefied to the minimum feature count (9,869). Pairwise comparisons of alpha diversity metrics showed no significant differences between aquifers within the same year, nor between 2021 and 2022 samples from the same aquifer (Kruskal-Wallis, p>0.05). Individual samples are colored by the total arsenic concentration measured at the well.

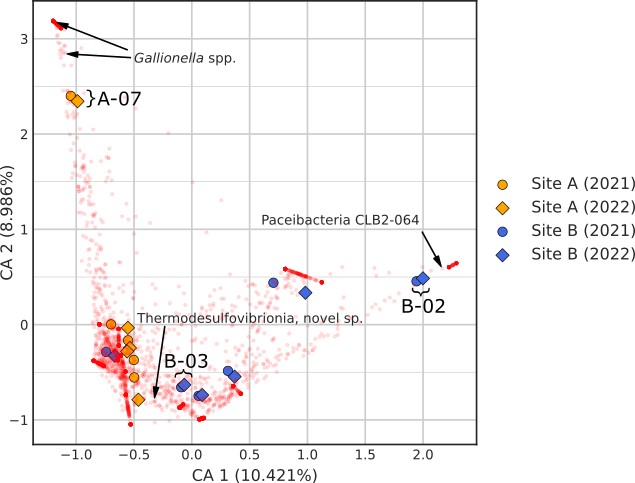

**Figure S2: Correspondence Analysis (CA) biplot.** Red points indicate the position of features (ASVs) with respect to the CA axes. Features of interest are annotated with arrows: a novel Paceibacteria ASV (corresponding to the metagenome-assembled genome Paceibacteria CLB2-064), high abundance in well B-02; a novel Thermodesulfovibrionia ASV, high abundance in well B-03; and various novel *Gallionella* ASVs, high abundance in well A-07. Samples of interest (A-07, B-02, B-03) associated with the indicated features are annotated with brackets.

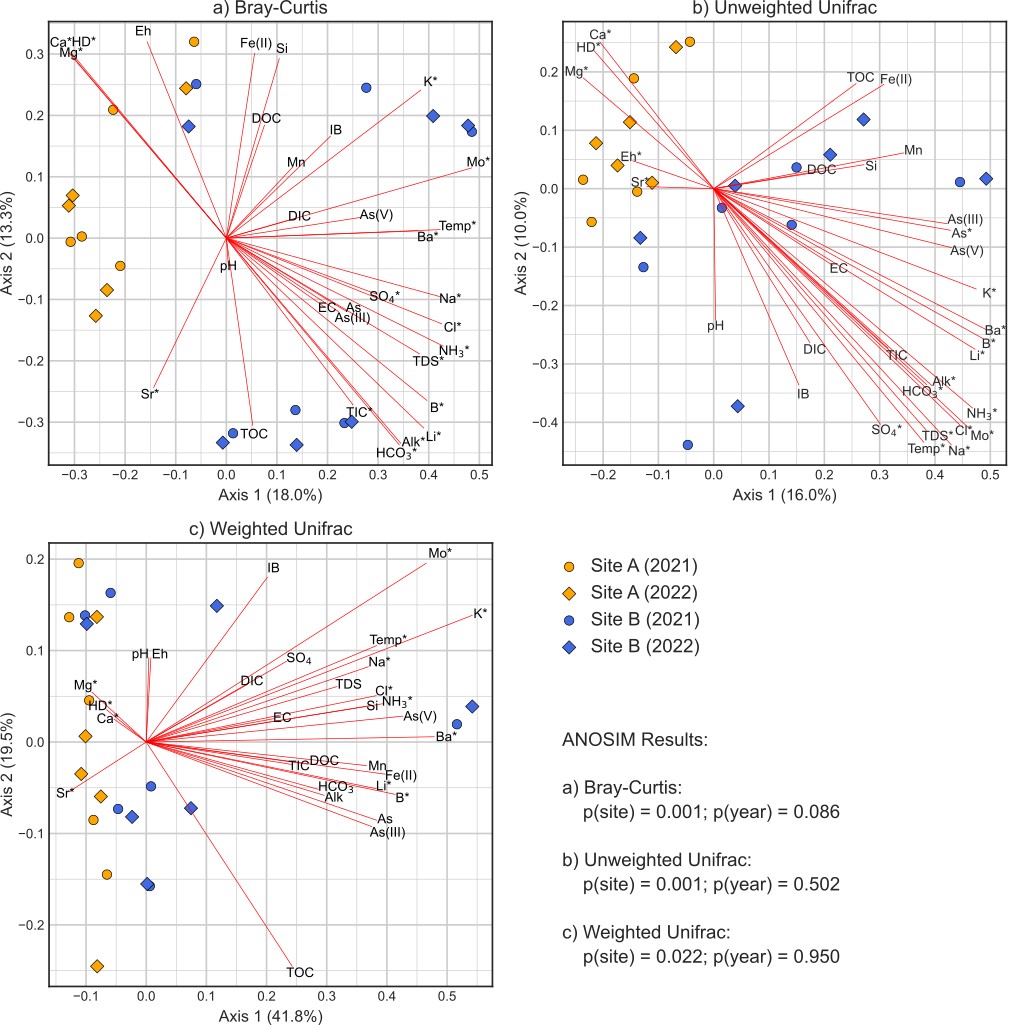

**Figure S3: Principal coordinates analysis (PCoA) plots** of a) Bray-Curtis, b) Unweighted Unifrac, and c) Weighted Unifrac distance matrices. Points represent samples, which are colored by site (aquifer), with samples taken in different years indicated by shape. Significant differences (p<0.05) were found between Sites A and B, but not between 2021 and 2022 samples (ANOSIM). Geochemical variables are projected on each ordination as loading vectors, with significant variables (p<0.05, see Table S1) indicated with an asterisk. Abbreviation key: EC = Electrical Conductivity, Temp = Temperature, As = Total Arsenic, DIC = Dissolved Inorganic Carbon, DOC = Dissolved Organic Carbon, HD = Hardness, IB = Ion Balance, Alk = Alkalinity, TDS = Total Dissolved Solids, TIC = Total Inorganic Carbon, TOC = Total Organic Carbon.

**Table S1: Groundwater geochemistry measurements from sampled wells.** Geochemical factors with measurable values in all wells in both 2021 and 2022, and which are predicted to influence arsenic biogeochemical cycling are included. Values are reported in mg/L unless otherwise indicated. The associated p-values from environmental fitting of factors to sample ordination scores (from PCoA; see Supplemental Figure 1) are reported in the bottom 3 rows; significant p-values (<0.05) are indicated with an asterisk.

| Sample | Year | pH | Eh (mV) | Temp. (°C) | Ammonia (as N) | Arsenic (total) | As (III) | As (V) | DOC | Fe (II) | Mn | SO_4_ | Alkalinity (as CaCO_3_) |
| --- | --- | --- | --- | --- | --- | --- | --- | --- | --- | --- | --- | --- | --- |
| A-01 | 2021 | 7.51 | -55.3 | 15.3 | 1.1 | 0.016 | 0.0145 | 0.00105 | 5.0 | 2.0 | 0.095 | 28 | 490 |
| A-02 | 2021 | 7.48 | -61.0 | 15.0 | 1.2 | 0.023 | 0.0165 | 0.00452 | 4.4 | 1.9 | 0.088 | 26 | 490 |
| A-03 | 2021 | 7.06 | -23.5 | 12.3 | 1.3 | 0.022 | 0.0195 | 0.00152 | 5.4 | 2.2 | 0.099 | 32 | 470 |
| A-04 | 2021 | 7.32 | -48.6 | 16.0 | 1.3 | 0.035 | 0.0307 | 0.00221 | 5.2 | 2.1 | 0.10 | 27 | 470 |
| A-07 | 2021 | 6.94 | 35.7 | 11.2 | 1.1 | 0.029 | 0.0239 | 0.00380 | 6.5 | 1.5 | 0.12 | 22 | 470 |
| B-02 | 2021 | 6.76 | -129.9 | 26.3 | 2.9 | 0.17 | 0.155 | 0.00966 | 5.5 | 4.0 | 0.20 | 38 | 530 |
| B-03 | 2021 | 7.23 | -133.3 | 16.3 | 2.8 | 0.25 | 0.243 | 0.0104 | 5.3 | 2.5 | 0.20 | 48 | 580 |
| B-04 | 2021 | 7.41 | -110.7 | 32.6 | 2.6 | 0.025 | 0.0202 | 0.00170 | 4.8 | 0.28 | 0.035 | 53 | 550 |
| B-05 | 2021 | 7.23 | -130.3 | 26.0 | 3.1 | 0.08 | 0.0736 | 0.00307 | 5.7 | 1.2 | 0.074 | 52 | 540 |
| B-06 | 2021 | 7.21 | -47.4 | 22.5 | 2.7 | 0.016 | 0.0121 | 0.00293 | 5.7 | 0.12 | 0.084 | 50 | 570 |
| B-07 | 2021 | 7.77 | -140.1 | 8.6 | 2.2 | 0.036 | 0.0327 | 0.00183 | 4.8 | 1.5 | 0.13 | 59 | 530 |
| A-01 | 2022 | 7.20 | -99.5 | 14.7 | 1.1 | 0.014 | 0.0169 | 0.00105 | 8.1 | 2.1 | 0.090 | 27 | 510 |
| A-02 | 2022 | 7.38 | -102.8 | 12.7 | 1.3 | 0.018 | 0.0196 | 0.00380 | 7.2 | 1.9 | 0.085 | 28 | 500 |
| A-03 | 2022 | 7.50 | 155.2 | 10.5 | 1.2 | 0.019 | 0.0243 | 0.00174 | 6.5 | 2.3 | 0.096 | 38 | 490 |
| A-04 | 2022 | 7.37 | 160.7 | 14.0 | 1.3 | 0.025 | 0.0367 | 0.00283 | 8.2 | 2.2 | 0.10 | 29 | 480 |
| A-07 | 2022 | 7.10 | 69.4 | 11.2 | 1.2 | 0.024 | 0.0289 | 0.00612 | 6.9 | 1.7 | 0.12 | 28 | 480 |
| B-02 | 2022 | 7.16 | -142.5 | 24.7 | 3.0 | 0.15 | 0.176 | 0.0135 | 8.6 | 4.2 | 0.20 | 44 | 540 |
| B-03 | 2022 | 7.45 | -150.3 | 15.0 | 3.3 | 0.24 | 0.289 | 0.0143 | 7.0 | 3.2 | 0.24 | 63 | 590 |
| B-04 | 2022 | 7.46 | -109.1 | 31.5 | 2.8 | 0.024 | 0.0266 | 0.00351 | 6.8 | 0.49 | 0.049 | 55 | 550 |
| B-05 | 2022 | 7.70 | -138.9 | 27.2 | 3.1 | 0.067 | 0.0687 | 0.00453 | 6.1 | 1.2 | 0.079 | 58 | 560 |
| B-06 | 2022 | 7.95 | 44.2 | 23.9 | 2.7 | 0.014 | 0.0126 | 0.00329 | 6.0 | 0.10 | 0.077 | 54 | 570 |
| B-07 | 2022 | 7.78 | -153.0 | 6.9 | 2.4 | 0.029 | 0.0355 | 0.00541 | 5.9 | 1.5 | 0.13 | 67 | 540 |
| p (Bray-Curtis) | | 0.918 | 0.134 | 0.001* | 0.001* | 0.06 | 0.085 | 0.125 | 0.975 | 0.736 | 0.822 | 0.011* | 0.001* |
| p (Unweighted Unifrac) | | 0.797 | 0.029* | 0.001* | 0.001* | 0.041* | 0.051 | 0.059 | 0.847 | 0.882 | 0.561 | 0.004* | 0.001* |
| p (Weighted Unifrac) | | 0.488 | 0.291 | 0.009* | 0.035* | 0.144 | 0.17 | 0.241 | 0.588 | 0.703 | 0.689 | 0.527 | 0.246 |

**Data File S1: Groundwater analysis report.** Additional information on the laboratory methods and measured variables from groundwater samples analyzed by Brooks Applied Labs (pages 1-2) and Bureau Veritas (pages 3-4).

**Data File S2:**

**Tab 1: All geochemical variables.** All measured values, with associated units, of groundwater samples associated with each sampling site.

**Tab 2: envfit results.** The output generated by environmental fitting (envfit) of geochemical variables to Bray-Curtis, unweighted unifrac, and weighted unifrac ordinations. Only variables that were measured in both 2021 and 2022, with values above the detection limit in all wells, were included. Significant correlations are indicated with asterisks (see Signif. codes). Note that PC1 and PC2 values do not necessarily represent the coordinates of loading vectors displayed on Figure S3, as vectors were scaled to the PCoA axes ranges prior to plotting.

**Tab 3: Metagenome-assembled genome (MAG) details.** Information on all high-quality MAGs is included. The first four characters of MAG ids (Bin) represent the site (CLA = Site A, CLB = Site B) and location/well of the sample (by number, e.g., CLB2 = Site B-02). Completeness and contamination scores were calculated by CheckM. Percent relative abundance is expressed as the average scaffold coverage from Bowtie2 read mapping within a MAG, divided by the sum of all average coverage values across high-quality bins within the sample.

**Tab 4: DRAM pathways.** The output table generated by DRAM, where metabolic genes are aggregated into pathways for each MAG. Pathways are described with either percentage completion values (0-100%) or as true/false.

**Tab 5: CLB7-064 proportionality.** Summary of all MAGs with scaffolds showing proportional abundance to CLB7-064 scaffolds**.** Scaffold (>10 kb) proportionality was calculated using the ‘rho’ metric in the propr R package, with a cut-off of 0.9 indicating significant proportional abundance. No unbinned scaffolds were significantly proportional to CLB7-064. Only one MAG (indicated with an asterisk) also showed significant proportionality to CLB7-064 when comparing MAG relative abundances instead of individual scaffolds. The % of genome proportional column identifies the percent of the total genome length made up of scaffolds scoring as proportionally abundant.
